# Phenomics Assistant: An Interface for LLM-based Biomedical Knowledge Graph Exploration

**DOI:** 10.1101/2024.01.31.578275

**Authors:** Shawn T O’Neil, Kevin Schaper, Glass Elsarboukh, Justin T Reese, Sierra A T Moxon, Nomi L Harris, Monica C Munoz-Torres, Peter N Robinson, Melissa A Haendel, Christopher J Mungall

**Affiliations:** Department of Biomedical Informatics, University of Colorado Anschutz Medical Campus, Aurora, CO, 80045, USA; Environmental Genomics and Systems Biology, Lawrence Berkeley National Laboratory, Berkeley, CA, 94720, USA; The Jackson Laboratory for Genomic Medicine, Farmington CT, 06032, USA

## Abstract

We introduce Phenomics Assistant, a prototype chat-based interface for querying the Monarch knowledge graph (KG), a comprehensive biomedical database. While unaided Large Large Language models (LLMs) are prone to mistakes in factual recall, their strong abilities in summarization and tool use suggest new opportunities to help non-expert users query and interact with complex data, while drawing on the KG to improve reliability of the answers. Leveraging the ability of LLMs to interpret queries in natural language, Phenomics Assistant enables a wide range of users to interactively discover relationships between diseases, genes, and phenotypes.

To assess the reliability of our approach and compare the accuracy of different LLMs, we evaluated Phenomics Assistant answers on benchmark tasks for gene-disease association and gene alias queries. While comparisons across tested LLMs revealed differences in their ability to interpret KG-provided information, we found that even basic KG access markedly boosts the reliability of standalone LLMs. By enabling users to pose queries in natural language and summarizing results in familiar terms, Phenomics Assistant represents a new approach for navigating the Monarch KG.

## Introduction

Large language models (LLMs) represent a new paradigm in human-computer interaction, allowing users to work with systems in their native language. LLMs excel in summarizing, paraphrasing, and explaining in-context information [1,2], spurring the growth of knowledge-backed AI agents, capable of using tools to search for and contextualize externally-sourced information [3]. The information they generate, however, is not always accurate, particularly for information that is not well represented in training data [4].

Knowledge graphs (KGs) are a powerful approach for integrating heterogeneous data and enabling the data to be queried to discover new insights; they are widely used in biomedicine and beyond. In translational research, KGs are frequently used to represent known relationships between biomedical entities such as diseases, genes, and phenotypes, where insights into these relationships can lead to improved treatments [5]. The Monarch Initiative KG includes millions of known, curated associations across hundreds of thousands of entities for dozens of species [6]. However, the sheer volume and complexity of genetic data pose significant challenges in terms of accessibility and interpretation. While there are a number of interfaces for querying such data (Figure 1), including specialized query languages [7], graphical interfaces [8], and information-rich websites and APIs [6,9], using these effectively often requires domain-specific vocabulary and knowledge, limiting their utility for a broader range of users, including clinicians and researchers without extensive bioinformatics training. Integrating the translational information stored in biomedical KGs with the user-friendly features of LLMs thus presents a promising direction for enhancing the accessibility of the Monarch KG.

**Figure 1:**
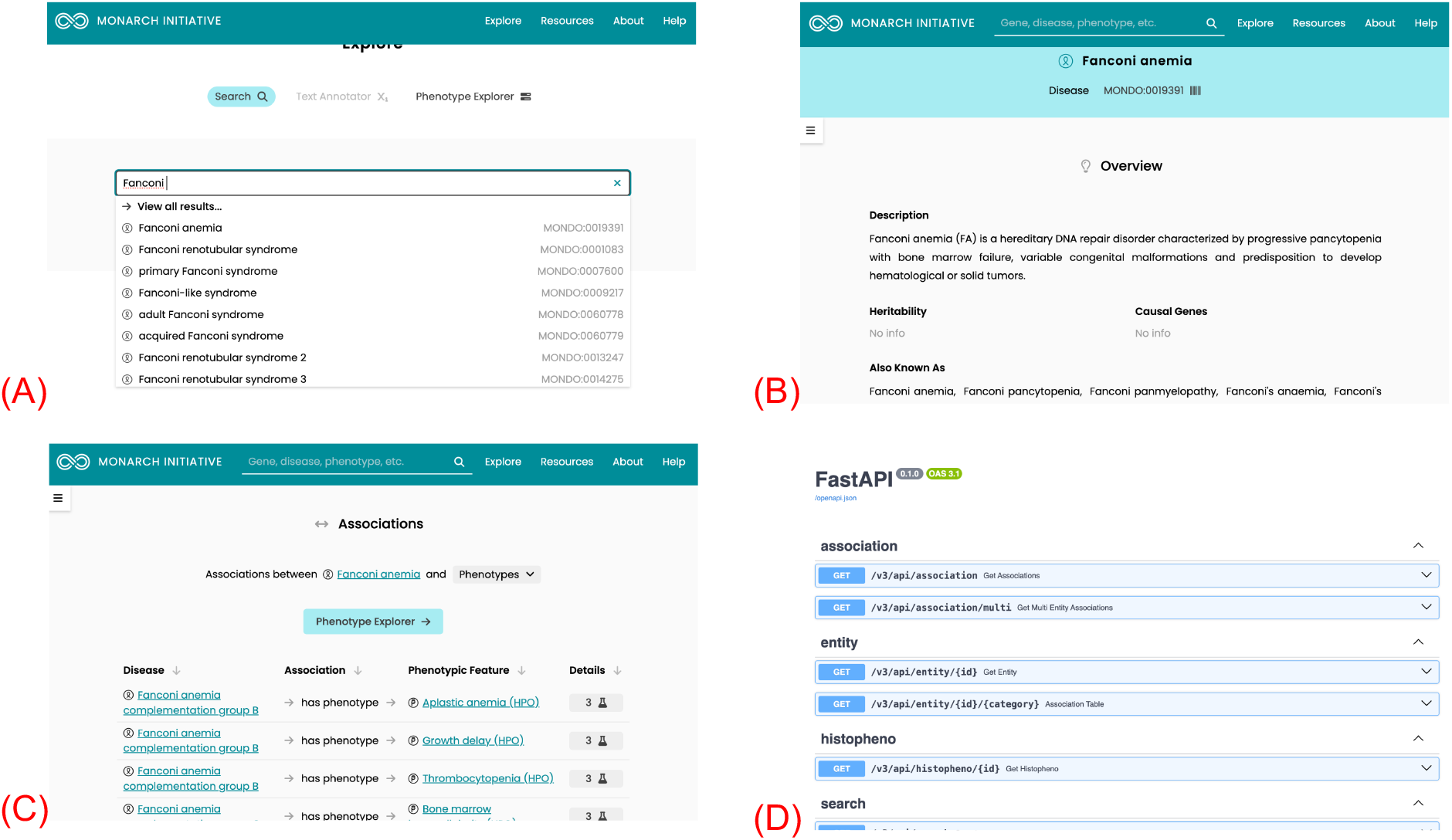
Monarch Initiative interfaces. While biomedical databases such as Monarch host vast amounts of information, their interfaces are generally designed for domain experts to search (A), quickly review related information (B) and known associations (C), or provide access via API (D).

In this paper we introduce Phenomics Assistant, an LLM-based interface for searching, retrieving, and summarizing information in the Monarch Initiative KG. We describe its prototype implementation as a user-friendly web application, and analyze its performance on natural language benchmark tasks for gene name identification and gene-disease associations, comparing several LLM models with and without KG access. These results highlight the importance of providing curated KG information to LLMs, and reveal differences in the ability of different LLMs to use the provided information. On average, LLMs with KG access produced between 1.9X and 5.1X more correct answers than those without KG access in our tests.

The integration of LLMs with domain-specific, curated knowledge bases like the Monarch Knowledge Graph presents a new avenue for scientific question-answering in the field of genomics. While LLMs are adept at generating coherent and contextually relevant responses, their reliance on training data can lead to inaccuracies or omissions, particularly in specialized domains [4]. Phenomics Assistant addresses this by grounding LLM responses in the verified data of the KG, enhancing the information provided with links to sources and other information. By allowing users to pose queries in natural language and summarizing data in familiar terms, Phenomics Assistant democratizes access to complex genomic information, making it more readily available to a diverse audience.

Phenomics Assistant is still in active development. A demonstration deployment can be accessed from the GitHub repository at https://github.com/monarch-initiative/phenomics-assistant.

### Related Work

Augmenting LLMs with external tools or curated data is a common approach to improving their factual accuracy and reasoning abilities [3]. Many LLM systems incorporate free-text document databases, for example to support question-and-answer tasks over scientific literature [10].

LLMs have demonstrated proficiency in accurately summarizing structured data, and they can be used with relational databases to assist non-specialists in data exploration and querying [11]. LLMs may also be configured to access APIs; this technique powers ChatGPT “plugins,” allowing the AI to access information or take action on users’ behalf [12], and has been used to access data via scientific APIs such as PubMed’s E-utils [13].

A variety of applications have integrated LLMs and knowledge graphs, including assisting in the development or curation of KGs by extracting entities and relationships from free text [14–18]. Conversely, a number of researchers have explored providing KG data to LLMs to improve the reliability of the answers that are generated [19]. In biomedical applications, some approaches utilize semantic similarity search via embedding vectors. Recent approaches include embedding search followed by neighborhood retrieval, filtering the associations by query similarity for LLM summarization [20], neighborhood filtering based on a query classification [21], additionally considering document collections [22], rewriting neighborhood descriptions in text [23], decomposing queries into logical constructs [24], fine-tuning model weights with additional KG-sourced training data [25], and developing specialized graph neural networks for improved reasoning [26].

As we discuss below, Phenomics Assistant accesses KG information via the Monarch API rather than directly, placing it closer in spirit to ChatGPT plugins and other API-accessing LLM utilities, and thus more readily deployable over existing infrastructure. As our evaluations show, this API-backed approach makes effective use of KG structure, including information about classes of diseases and phenotypes, resulting in significant improvements over unaided LLMs.

## Methods

### Architecture

Phenomics Assistant consists of several components illustrated in Figure 2. Users interact with a prototype chat-based user interface that connects to an LLM framework, mediating access to a subset of functionality of the Monarch Initiative API.

**Figure 2:**
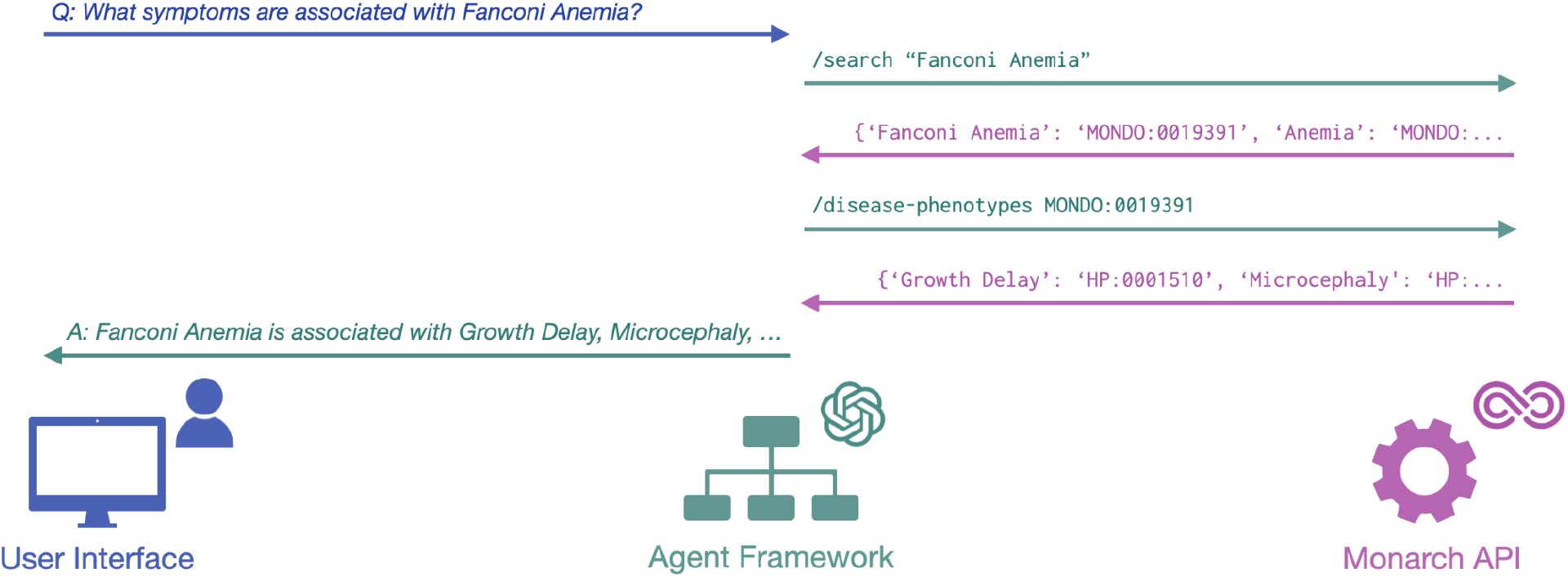
Phenomics Assistant architecture. Users pose questions via the UI in natural language (blue), and these are translated to Monarch API calls by the LLM and agent framework (green). Responses from the API (pink) are evaluated by the LLM, and may trigger followup calls and responses, until a final answer is returned to the user in natural language.

#### API

The standard Monarch Initiative API provides a wide variety of functions, including keyword-based entity search, flexible entity-association retrieval, and semantic similarity search. While LLMs can be adapted to call such functions as “tools,” minimizing the number and complexity of available tools reduces potential for errors caused by inappropriate tool use [27]. For Phenomics Assistant we thus utilize a small set of LLM-focused functions, including search and individual association-type lookups, described in Table 1. Importantly, the association endpoints consider subclass closures for queried entities, matching the behavior of the Monarch website interface. Gene associations for Ehler-Danlos Syndrome (MONDO:0017314), for example, will also include associations for the autosomal dominant and recessive subtypes (MONDO:0007524, MONDO:0002014). Gene associations always include both causal and correlated relationships, distinguishing between the two. Returned lists of associations include URLs to publications or evidence information when available from the KG.

**Table 1:**
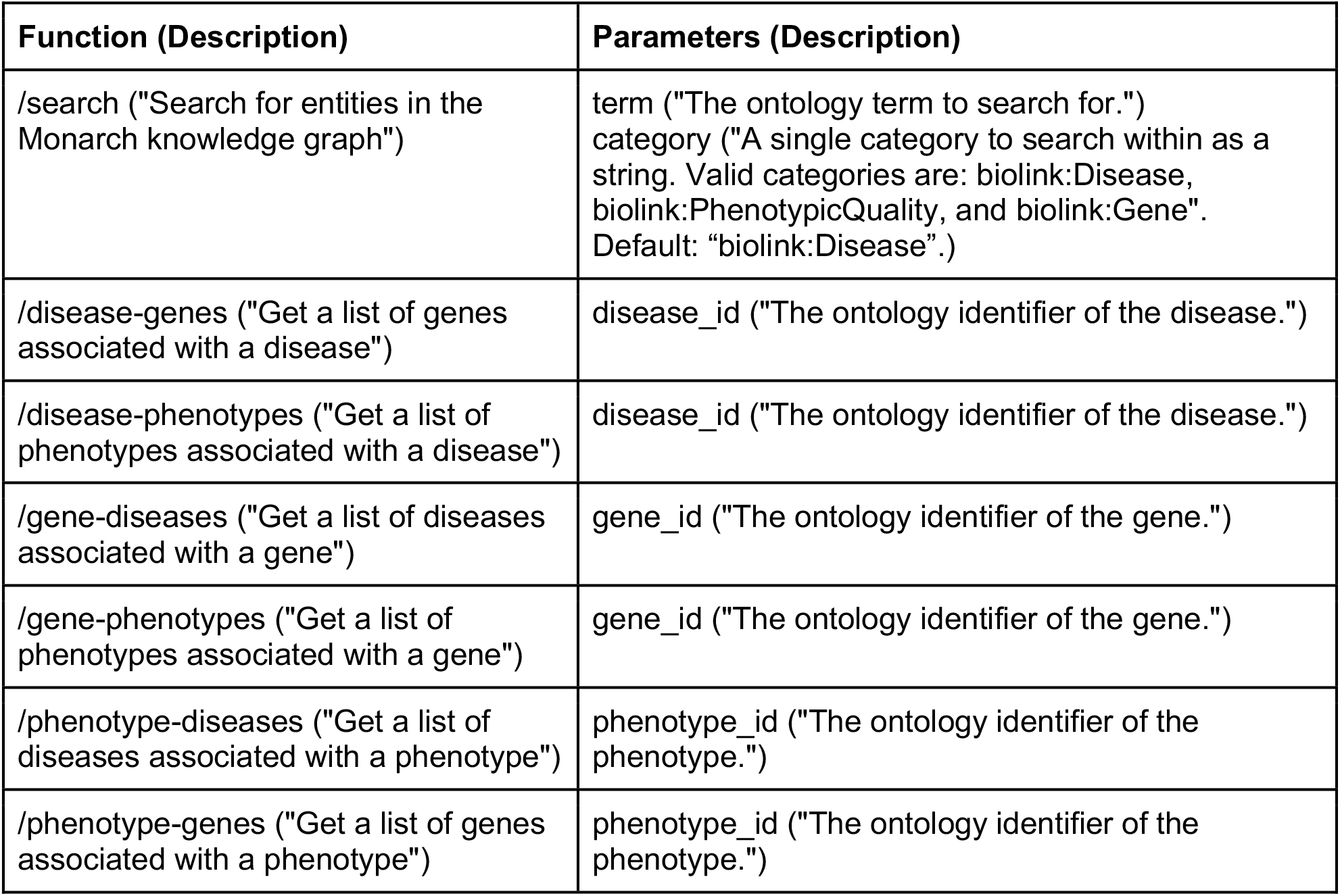
Available API functions and parameters. Function names, parameters, and their descriptions are supplied as part of the prompt to the LLM. All functions also include optional *limit* and *offset* parameters (described as part of the LLM prompt as “The maximum number of search results to return” and “Offset for pagination of results” respectively, with defaults 0 and 10).

#### Agent Framework

Some LLMs are trained to “call” external functions when provided with callable function metadata as part of the conversation context, by responding to queries with the names and parameters of functions to call. These specially-formatted responses are then parsed and executed locally before including the results in followup responses to the model. Utilizing OpenAI models that support function-calling [28], we developed an agent-based framework that extracts function metadata (functions, parameters, and descriptions from Table 1) and provides it to the LLM, executes LLM-specified calls, and returns results in JSON format. The framework also manages LLM choice (e.g. GPT 3.5 or GPT 4), conversation history, toxicity checks using OpenAI’s moderation API [29], and the system prompt, which is used by many LLMs to guide model behavior. Phenomics Assistant’s system prompt instructs the model to use lay language descriptions and include links to external pages when possible (Suppl. Table 1).

#### User Interface

A web application provides a chat interface to Phenomics Assistant. The current interface is developed with the Streamlit web framework, enabling rapid feature prototyping. Users enter questions in natural language, and the interface provides real-time feedback on functions being called, optionally including call parameters and results in the conversation stream (Figure 3). Responses include formatted links to external pages or publications when available (Figure 4). Such transparency features can increase confidence in results by allowing users to check answers against returned data and follow up with external resources.

**Figure 3:**
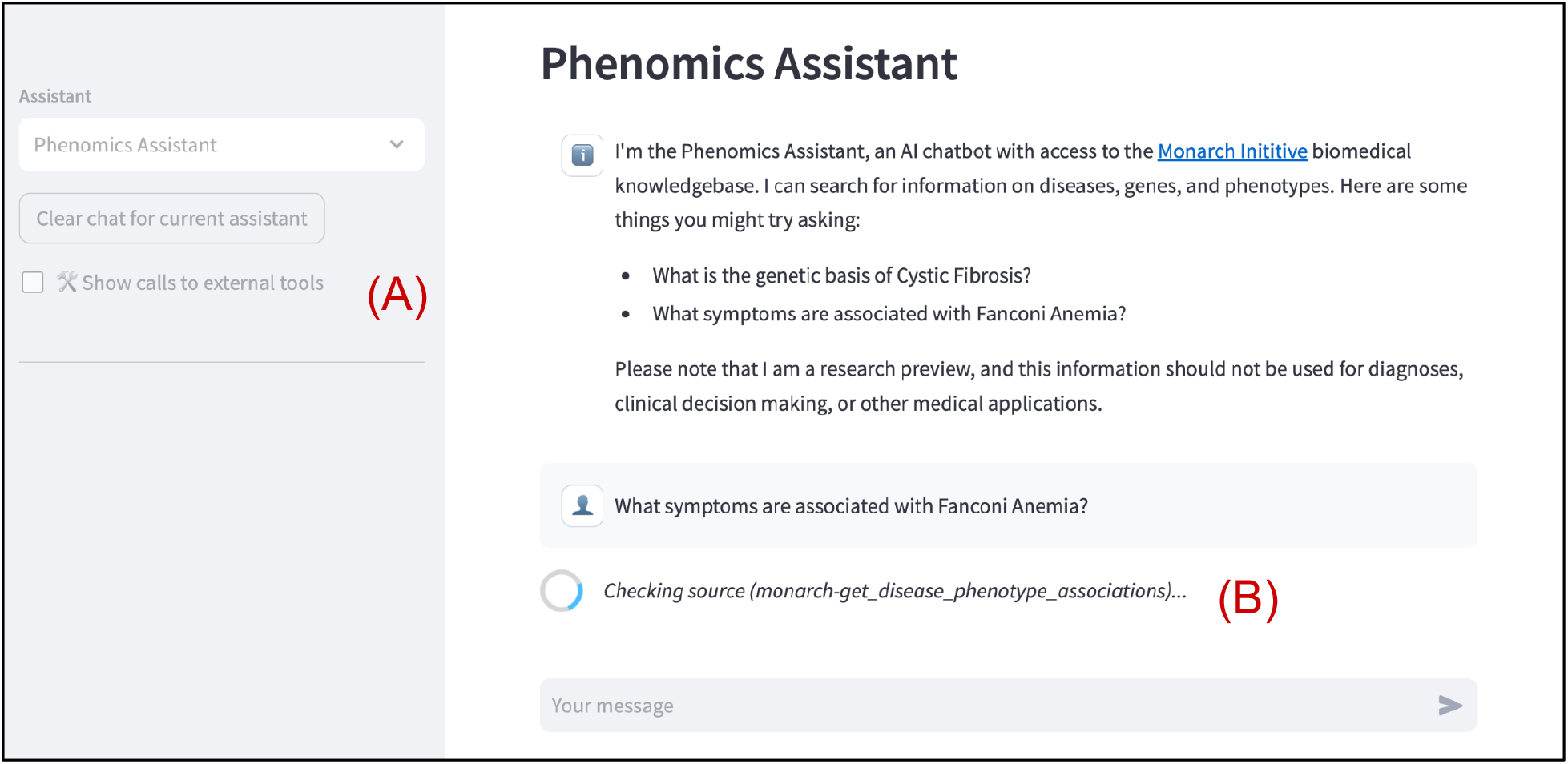
Phenomics Assistant prototype user interface. Users can select from different Assistant variations and optionally include calls and responses in the conversation stream (A). Real-time feedback alerts the user to specific endpoints or functions being called (B).

**Figure 4:**
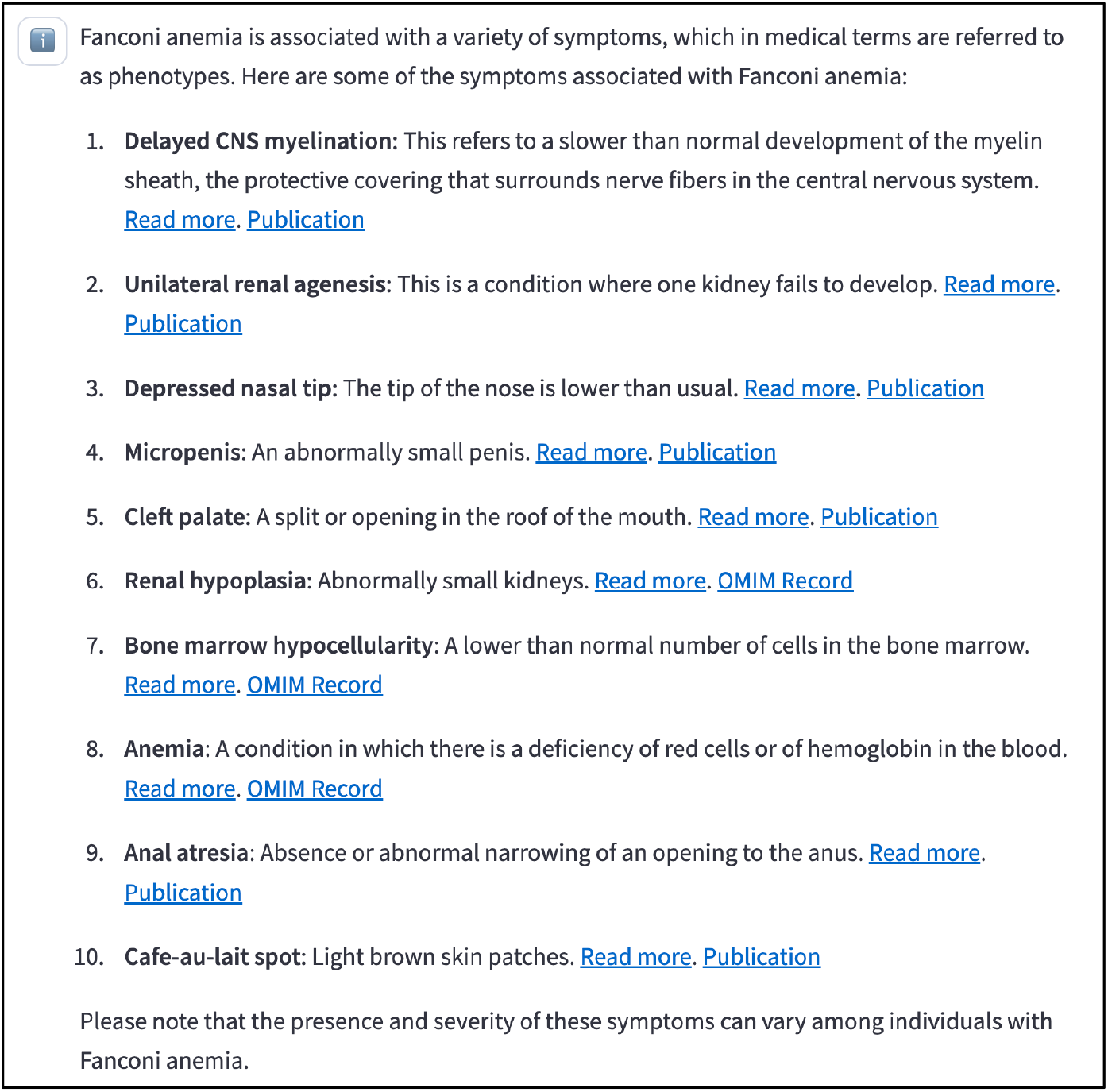
Example query response. Links are included to relevant pages and data sources when available. This response required two API calls (not shown), first to (`/search(` for “Fanconi Anemia” to retrieve the relevant entity identifier in the KG, and second to (`/disease-phenotypes(` to retrieve the relevant associations.

### Evaluation

We compared the accuracy of three different LLMs (OpenAI’s `gpt-3.5-turbo-0613, gpt-4-0613`, and `gpt-4-1106-preview` models) with and without KG access on the *gene alias* and *gene-disease association* tasks of the GeneTuring benchmark dataset [30]. Each task consists of 50 question and gold-standard answer pairs. Gene alias questions ask for an official gene symbol for a non-standard gene name, for example “What is the official gene symbol of LMP10?” with the gold-standard answer being “PSMB10”. For this task, the GeneTuring authors specify a Jaccard similarity score, comparing the set of symbols mentioned in a given answer to the size-one set containing the gold standard, thereby penalizing additional mentioned names. Gene-disease association asks for the set of gene names associated with a disease, for example “What are genes related to Distal renal tubular acidosis?” with the gold-standard answer being “SLC4A1, APT6V0A4”. This task prescribes a recall metric, computed as the percentage of gold-standard genes mentioned in an answer (thus not penalizing additional mentioned names). The GeneTuring authors sourced data for the gene alias task from NCBI, and for gene-disease associations from OMIM [31], which is also a source for Monarch data[6].

All models use `temperature = 0`, minimizing (but not eliminating) stochastic variation in model responses. Models with KG access are configured similarly to those available in the Phenomics Assistant web interface at the time of testing, including the system prompt (Suppl. Table 1). Models without KG access were given the system prompt “You are a helpful assistant.”

Agent answers are provided as free text in markdown format (Figure 4). Computing metric scores thus requires extracting identifiers mentioned in answers for comparison to gold-standard answers. For example, given the question “What are genes related to Congenital disorder of deglycosylation?” with expected gold-standard answer of “MAN2C1, NGLY1” and LLM-generated answer of “Genes associated with Congenital disorder of deglycosylation include MAN2C1, PMM2, and ALG6”, we must compute the recall of the set (MAN2C1, PMM2, ALG6) against (MAN2C1, NGLY1). We accomplish this in an automated fashion by instantiating `gpt-4-0613`-based “evaluator” agents provided with functions that compute recall or Jaccard scores as appropriate from generated answer text. These LLM-callable functions take entity lists as parameters and return computed scores, allowing the evaluators to use function-calling for simultaneous named entity extraction and computational evaluation. While powerful, such model-based evaluations themselves require validation [32], particularly when applied to model-supplied outputs [33]. To validate this approach, we compute scores manually for 20 questions selected at random for each task (40 total) for comparison to evaluator-produced scores. Finally, we use single-sided, paired Wilcoxon signed-rank tests (`wilcox.test` in R version 4.2.2) to assess improvement in scores for models with KG access compared to those without.

## Results

The gene alias and disease-association tasks prescribe Jaccard similarity and recall metrics respectively, with both having a [0,1] range and 1.0 perfect score. Across models and tasks our results were largely bimodal, with most answers scoring 0.0 or 1.0. Manual evaluation for 20 random questions (out of 300 across tested models) from each task identified between 0 and 6 gene names per answer for scoring; LLM-evaluator extractions and scores agreed in 100% of these questions using case-sensitive exact-match criteria.

Figure 5 illustrates score counts across agents and tasks, revealing strong performance increases for models that are able to query the knowledge graph. For gene-disease associations, addition of knowledge graph resources increased the number of fully correct (score 1.0) answers by an average of 1.9X across models, and decreased wholly incorrect (0.0) answers by an average of 5.3X. Correct answer counts for gene alias increased by 5.1X with KG access and incorrect answer counts decreased by 2.7X. Table 2 analyzes these gains statistically; with the exception of GPT 3.5 for the gene-diseases association task, the estimated median score improvement with KG access is 1.0.

**Table 2:** Statistical comparisons of answer scores with and without KG access, per base model and task. Scores and score differences are non-normal, so we use single-sided paired Wilcoxon signed rank tests to estimate median score improvements for models with KG access compared to those with no access. *p* values are Bonferroni corrected across the six tests.

| Task | Base model | Estimated median KG-access score improvement | *p* value (adjusted) |
| --- | --- | --- | --- |
| Gene alias | GPT 3.5 | 1.0 | 0.00000482 |
| Gene alias | GPT 4 | 1.0 | 0.0000113 |
| Gene alias | GPT 4-1106 | 1.0 | 0.00000158 |
| Gene disease association | GPT 3.5 | 0.75 | 0.000167 |
| Gene disease association | GPT 4 | 1.0 | 0.0000158 |
| Gene disease association | GPT 4-1106 | 1.0 | 0.000193 |

**Figure 5:**
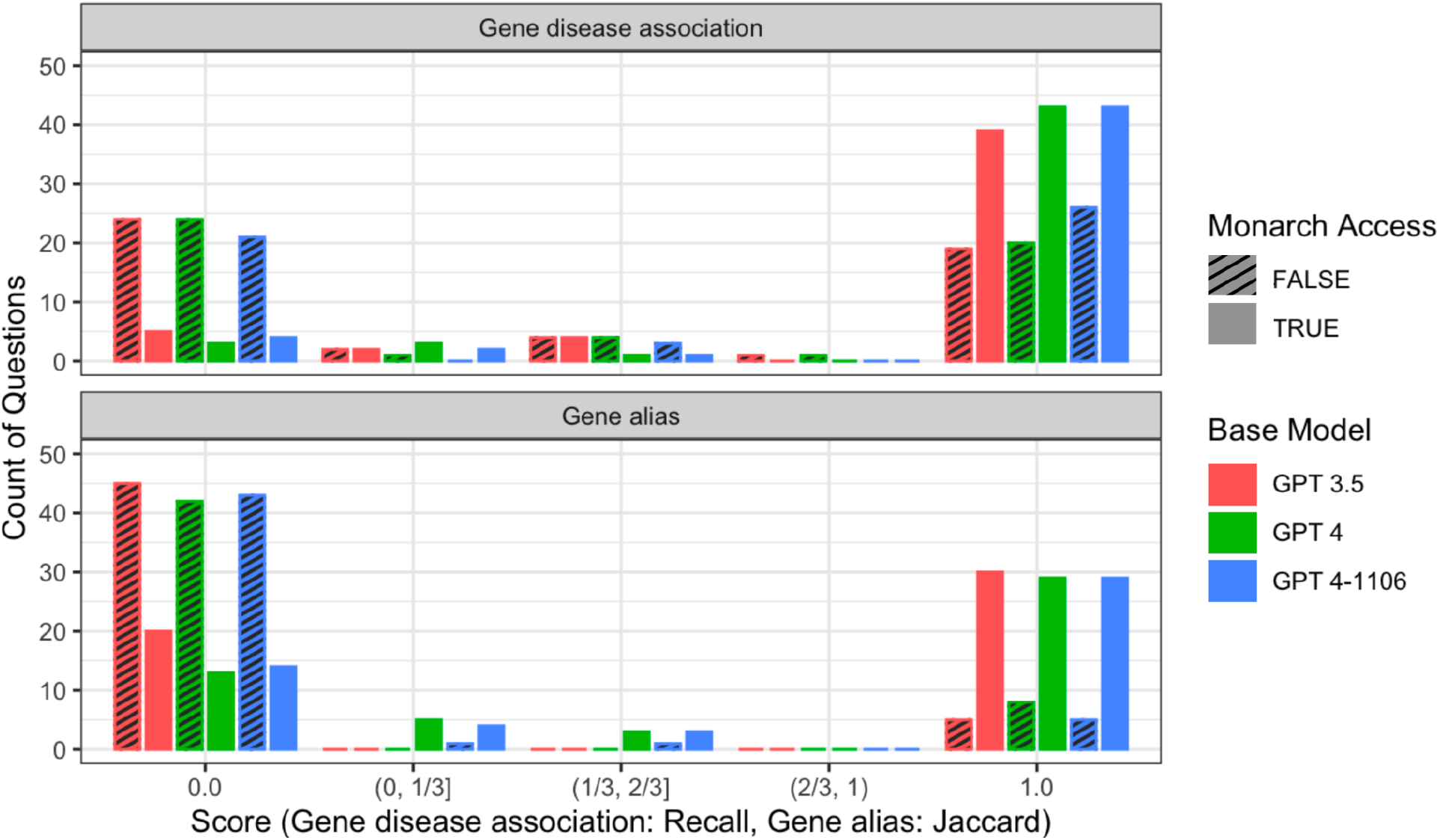
Evaluation results. Results are shown for three models (`gpt-3.5-turbo-0613, gpt-4-0613`, and `gpt-4-1106-preview`) with and without access to the Monarch knowledge graph API, on two GeneTuring evaluation tasks, gene-disease association and gene alias. Scores of 0.0 and 1.0 are counted separately; other scores are binned according to ranges shown in parentheses below the X axis.

Score differences across models with KG backing were small in comparison, suggesting that KG access is the major determinant of performance. For comparison, non-KG-backed GPT-3.5 scored 1.0 on 39% (19/50) of gene-disease answers, in line with results reported by the GeneTuring authors. The latest model, `gpt-4-1106-preview`, improves on this metric to 52% (26/50) without KG access.

### Model Comparisons

Although performance of GPT 3.5 and GPT 4-1106 was largely similar across the 100 posed questions, Table 3 lists 10 cases where the latter outperformed the former, and 4 cases of the opposite. Instances where GPT 4-1106 outperformed 3.5 appear to be influenced by suboptimal ordering of initial results from the /search function.. For example, a search for CXorf40B in response to “What is the official symbol of CXorf40B?” returns two entries: first CXorf40B, the name for chromosome X open reading frame 40B in the species *Gallus gallus*, and second EOLA2, the name for the homologous gene in *Homo sapiens*. Arguably, in this instance GPT 4-1106 better interpreted the question intent. Similarly, answering gene-disease questions requires a two step process: first, a search by disease name for the appropriate identifier, which may return multiple hits, followed by fetching associations for one of the results. In response to “What are genes related to Trichoepithelioma?” a search for “trichoepithelioma” returns multiple results, including Vulvar Trichoepithelioma (MONDO:0002201) first, followed by Familial Multiple Trichoepithelioma (MONDO:0011114). In this example GPT 3.5 followed up on the former, which is not associated with the prescribed gold-standard answer of CYLD, while 4-1106 followed up on the latter, which is. Similarly, a search for “proteasome-associated autoinflammatory syndrome” (PRAAS) matches multiple entries, and in the results PRAAS Type 5 (PRAAS5) is listed first; again, GPT 3.5 followed up with the first entry while 4-1106 chose the more appropriate generic condition (PRAAS) to fetch associations. As a final example, spinal muscular atrophy with congenital bone fractures (SMABF) has two types caused by different genes: SMABF1 caused by TRIP4, and SMABF2 caused by ASCC1. The search result listed both diseases and their descriptions, which include the causal gene names. Both 3.5 and 4-1106 followed up by fetching associations for the first, SMABF1. However, while 3.5 listed only the result of the latter query (TRIP4), 4-1106 included both gene names from the earlier search-provided information.

**Table 3:**
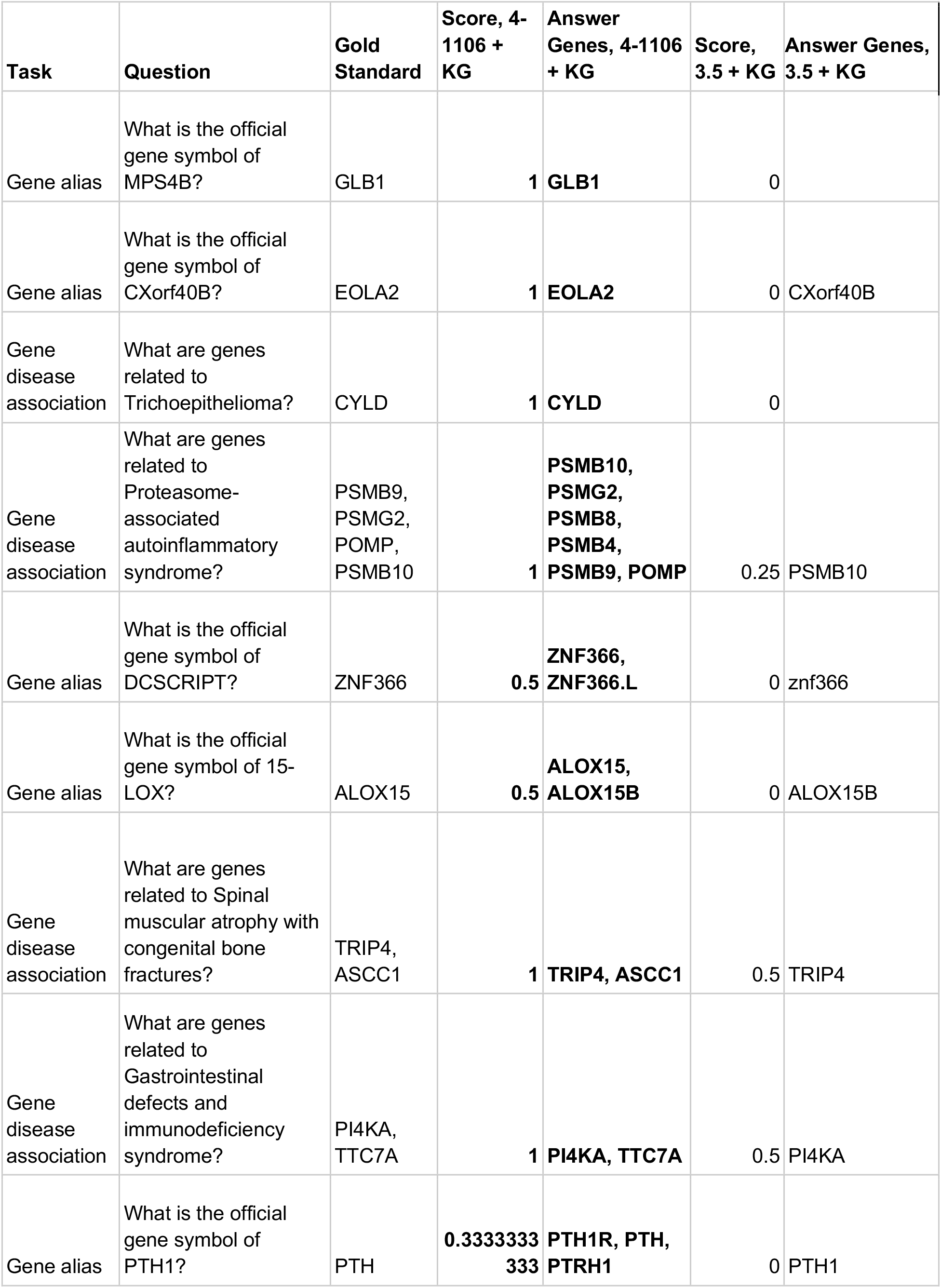
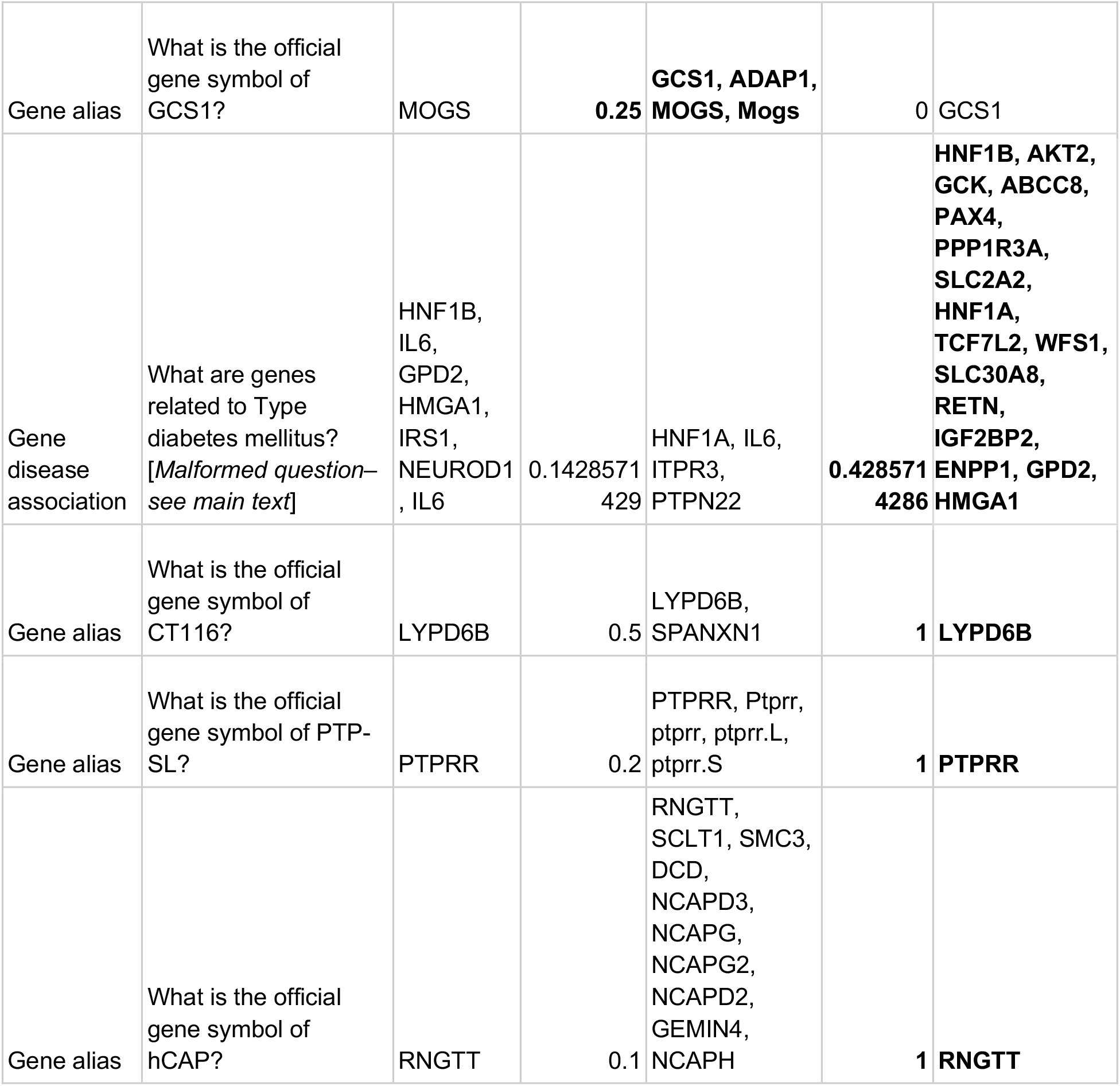
Questions for which GPT 4-1106 + KG performed differently than GPT 3.5 + KG. Better performance by GPT 4-1106 is largely due to improved interpretation of search results independent of result order. The best answers and scores in each row are bolded.

Table 3 lists four questions where GPT 3.5 outperformed 4-1106. Three of these are for gene alias questions, where GPT 4 included all results returned by the initial search, regardless of the species, while 3.5 listed only the first. The use of Jaccard similarity thus penalizes 4-1106’s comprehensiveness in comparison to 3.5. The question about diabetes is malformed - “What are genes related to Type diabetes mellitus?”. In this case the initial search listed Type I diabetes first and Type II diabetes second. GPT 3.5 followed up with associations for Type I, while GPT 4-1106 followed up with Type II.

Even for the most advanced GPT 4-1106 model, KG access resulted in increased scores on 30 gene alias and 19 gene-disease association questions (Suppl. Table 2). This finding is unsurprising, because LLMs are trained on large corpora of texts which may include inaccuracies, or may state factual truths in ways that are hard for a machine to understand. Of these 49 questions, GPT 4-1106 without KG access listed gene identifiers for 29, only three of which were partially correct (score > 0). GPT 4-1106 scored higher without KG access than with for only three questions; two of these were gene alias questions (seeking official gene names for DCSCRIPT and 15-LOX, see also Table 3) where KG-provided information provided additional context penalized by the Jaccard metric. The third sought genes related to Hyperphenylalaninemia. Here base GPT 4-1106 identified the six-gene gold standard set exactly, but when provided with KG access followed up on the first search result, Mild hyperphenylalaninemia (MONDO:0019335), which is only associated with one of these six.

Table 4 shows mean scores across questions per task for GPT 4-1106 with KG access, compared to similarly computed averages from other published methods on the same tasks. In the original GeneTuring work, Hou et al. consider multiple models, including GPT 3, ChatGPT (GPT 3.5), and New Bing (a version of GPT 4 with web search capabilities) [30]. Jin et al. developed GeneGPT, an approach similar to Phenomics Assistant with access to NCBI APIs and backed by the now discontinued OpenAI Codex model [13]. GPT 4-1106 with KG access performs best on gene-disease association tasks, slightly ahead of the web-enabled New Bing. GeneGPT performs best on gene alias tasks.

**Table 4:** **Mean task scores for 4-1106 + Monarch** compared to results reported by Hou et al. and Jin et al. [13,30].

| Task | GPT 3 | ChatGPT | New Bing | GeneGPT (best) | 4-1106 with KG |
| --- | --- | --- | --- | --- | --- |
| Gene disease association | 0.34 | 0.31 | 0.84 | 0.76 | **0.87** |
| Gene alias | 0.09 | 0.07 | 0.66 | **0.84** | 0.62 |

## Discussion

Phenomics Assistant is designed to help users to explore the primary entities in the Monarch KG – genes, diseases, and phenotypes – and known relationships between them. While the Monarch API provides keyword search and relationship information, the Assistant handles organization and summarization of the results for the user, along with links to available evidence and the Monarch website for more information. While not shown by default, the raw calls and retrieved data may be inspected by the user for transparency. In use, we have observed that the Assistant also successfully interprets incorrect or loosely-specified requests, for example searching for “Cystic Fibrosis” in response to “What genes are involved with CF?” and correcting misspelled disease names, such as searching for the correct “Ehlers-Danlos Syndrome” when asked about the incorrect “Ehlen-Danlos Syndrome.” On the other hand, while the natural-language descriptions of complex diseases and phenotypes are more accessible to a lay audience, they are in many cases generated by the LLM. While plausible, in a separate study we have observed that nuanced issues can infiltrate AI-generated definitions [15], and further work is needed to inform responses with KG-provided definitions.

Overall, we found that evaluation scores were significantly higher for models with access to Monarch data. For gene-disease associations the weakest model with KG access outperformed the strongest base model with 1.5X more correct answers; for gene aliases, over 3X more. However, unaided models performed reasonably for the gene-disease association task, with the latest GPT 4-1106 improving over GPT 3.5 with an additional 7 correct answers.

While access to Monarch data was the largest determinant of performance, model quality is also an important factor in how that data is utilized. This was particularly true when a list of retrieved information was given in suboptimal order. In several cases, GPT 3.5 used only the first entry returned, even when other choices would be more appropriate from the question context. Such order effects are well known [34], and can even contribute to hallucination (inaccurate results) in the presence of biased few-shot learning examples [35]. GPT 4-1106 was improved in this regard, and in one example (SMABF) incorporated information from both an initial search and subsequent association.

Finally, these results highlight the importance of both data representation and evaluation criteria when assessing performance of retrieval-based systems. All variants of Phenomics Assistant performed better on finding gene-disease associations than on identifying gene aliases, which may be expected given that source data for the former (OMIM) was present in the Monarch KG at the time of testing, while the source for the latter (NCBI gene aliases) was not. This distinction is reflected in opposite trends for GeneGPT, which utilizes NCBI APIs but not OMIM. This is confounded, however, by the different scoring criteria prescribed by the GeneTuring evaluation tasks. Gene-disease associations are scored according to the percentage of gold-standard genes identified, accommodating extra information provided in answers, whereas the Jaccard-based scoring for gene alias tasks does not. We prompted models to be comprehensive in answers, and since the Monarch KG contains information on many species, gene alias answers were frequently penalized by the Jaccard metric for providing aliases for multiple species, whereas GeneTuring references only human genes and diseases. Still, the same trend was seen for unaided base models, agreeing with the original GeneTuring test results and potentially impacted by GPT 4’s relative verbosity [36].

While Phenomics Assistant demonstrates good performance generally, open questions and future considerations remain. Many LLM-backed applications perform semantically-aware search via text embeddings [37], whereas the search functionality provided by the Monarch API is keyword-based. Given the observed sensitivity of some models to search result order, re-ranking results by embedding similarity, or implementing embedding-based semantic search directly, may further improve results. Similarly, we’ve implemented association retrieval between genes, diseases, and phenotypes, but the Monarch KG contains many other entity types and associations, and validating generic association-retrieval functionality for LLM use is a high priority.

Although we’ve focused the prototype Phenomics Assistant on the Monarch KG, additional access to other knowledge graphs could readily expand its capabilities to new domains. Furthermore, interface elements employed by many KG-driven applications such as plots, tables, and other widgets may complement the chat-only UI. Finally, we have thus far only tested OpenAI function-calling LLMs. Other closed-source models such as Claude (Anthropic) and Gemini (Google) have also started supporting function calling; some open source models have recently begun incorporating this feature [38].

Regardless of the base model, these results highlight the utility of LLM-based user interfaces in interacting with curated knowledge, dramatically improving the completeness and accuracy of generative AI models. Simultaneously, LLMs show promise in democratizing access to large, complex knowledge bases, effectively searching, summarizing, and contextualizing the information for end users.

## Data and Code Availability

Phenomics Assistant components are all open source and available on GitHub.

Main code repository: https://github.com/monarch-initiative/phenomics-assistant

User interface, with link to demonstration deployment: https://github.com/monarch-initiative/phenomics-assistant (see link in README)

API: https://github.com/monarch-initiative/oai-monarch-plugin

Agent framework: https://github.com/monarch-initiative/agent-smith-ai

Evaluation and results: https://github.com/monarch-initiative/oai-plugin-evals

## Supplementary Materials

**Suppl. Table 1:** System prompts used by models for evaluation. Information regarding the ‘function call setting’ pertains to functionality disabled during evaluation.

| Models | System Prompt |
| --- | --- |
| 3.5 + Monarch,<br>4 + Monarch,<br>4-1106 +<br>Monarch | You are the Monarch Assistant, an AI-powered chatbot that can answer questions about data from the Monarch Initiative knowledge graph.<br>You can search for entities such as genes, diseases, and phenotypes by name to get the associated ontology identifier.<br>You can retrieve associations between entities via their identifiers.<br>Users may use synonyms such as 'illness' or 'symptom'. Do not assume the user is familiar with biomedical terminology.<br>Always add additional information such as lay descriptions of phenotypes.<br>If the user changes the show function call setting, do not make any further function calls immediately.<br>IMPORTANT: Include markdown-formatted links to the Monarch Initiative for all results using the templates provided by function call responses. |
| 3.5 Base, 4<br>Base, 4-1106<br>Base | You are a helpful assistant. |

**Suppl. Table 2:**
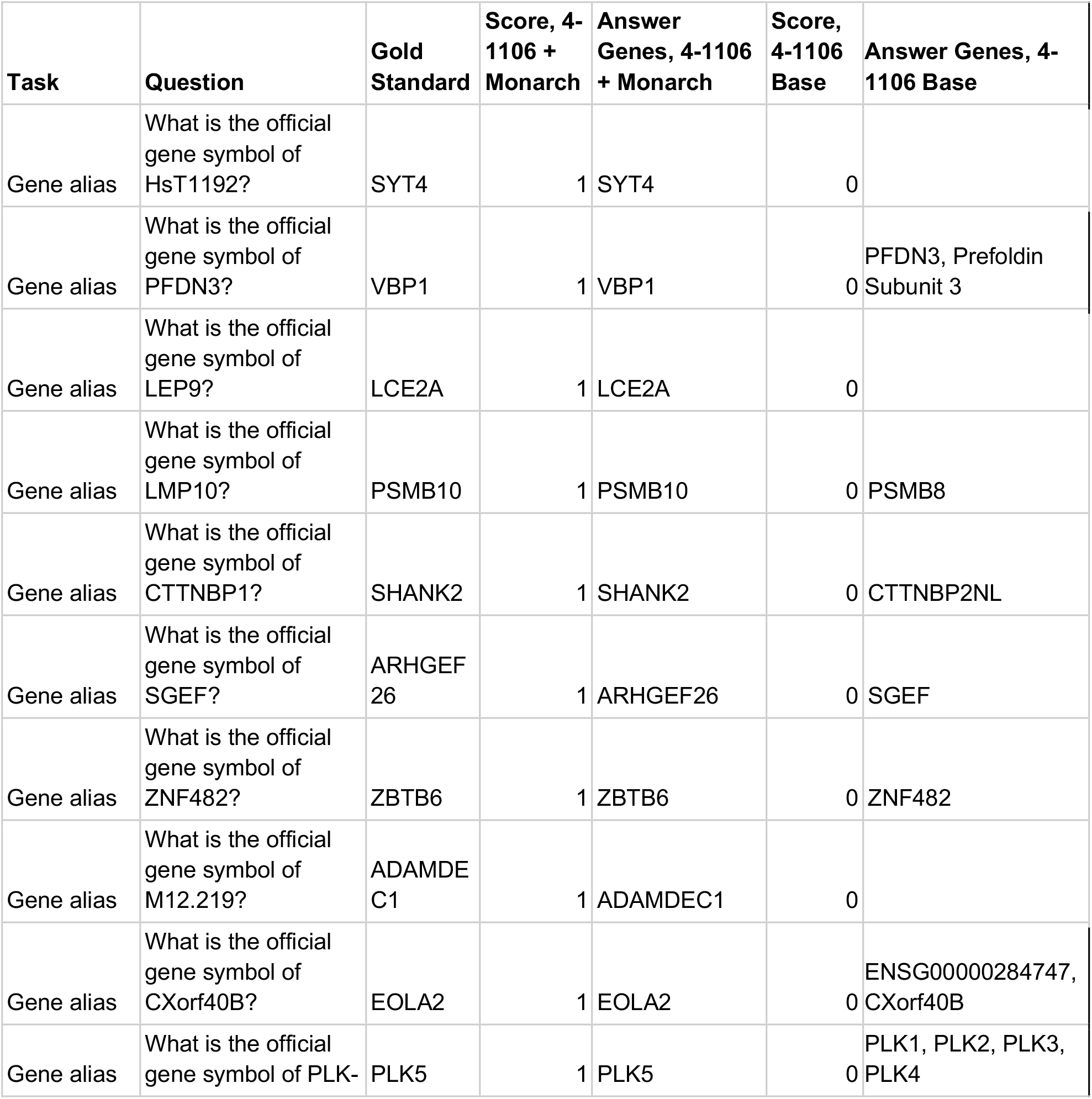
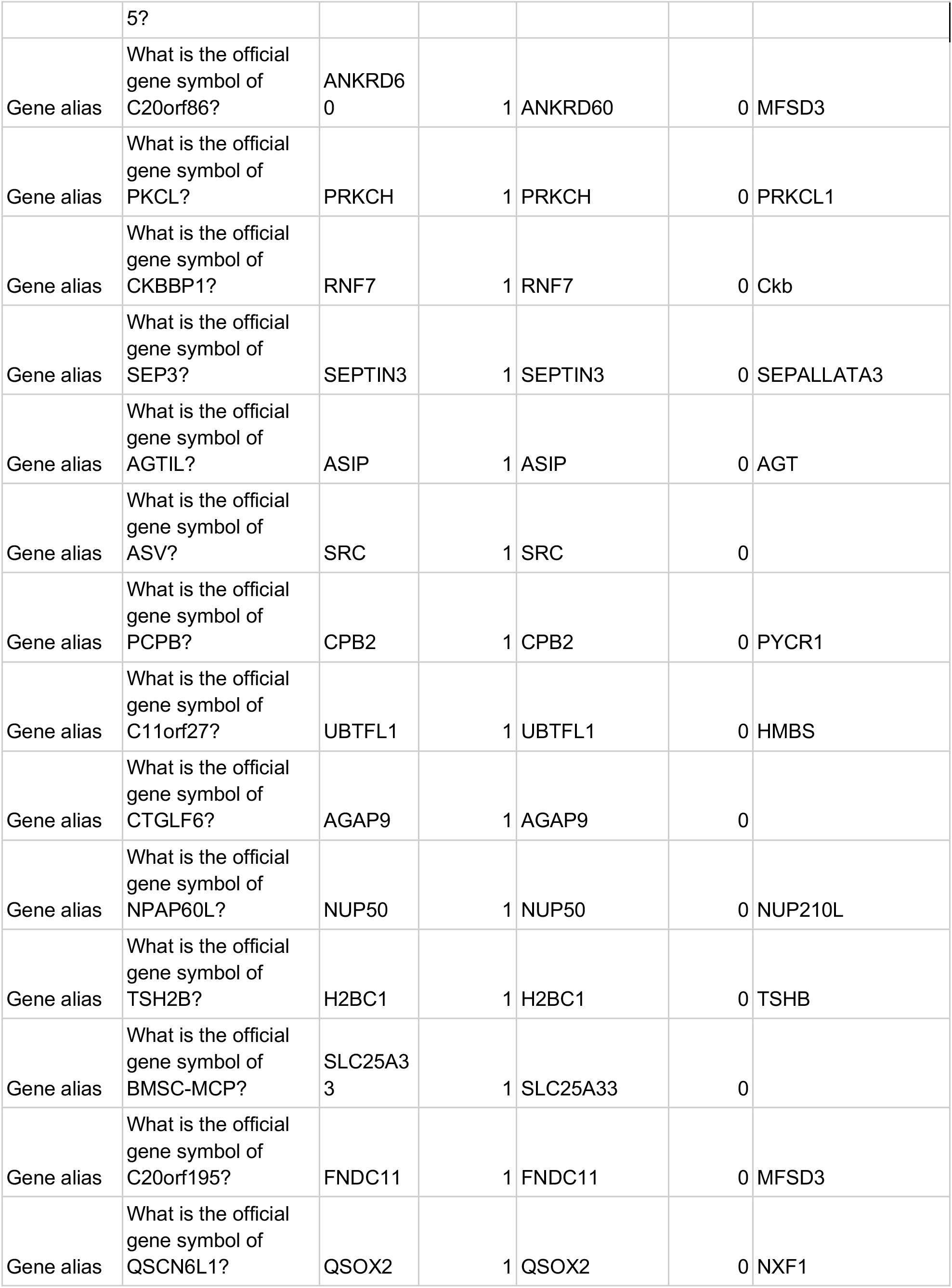
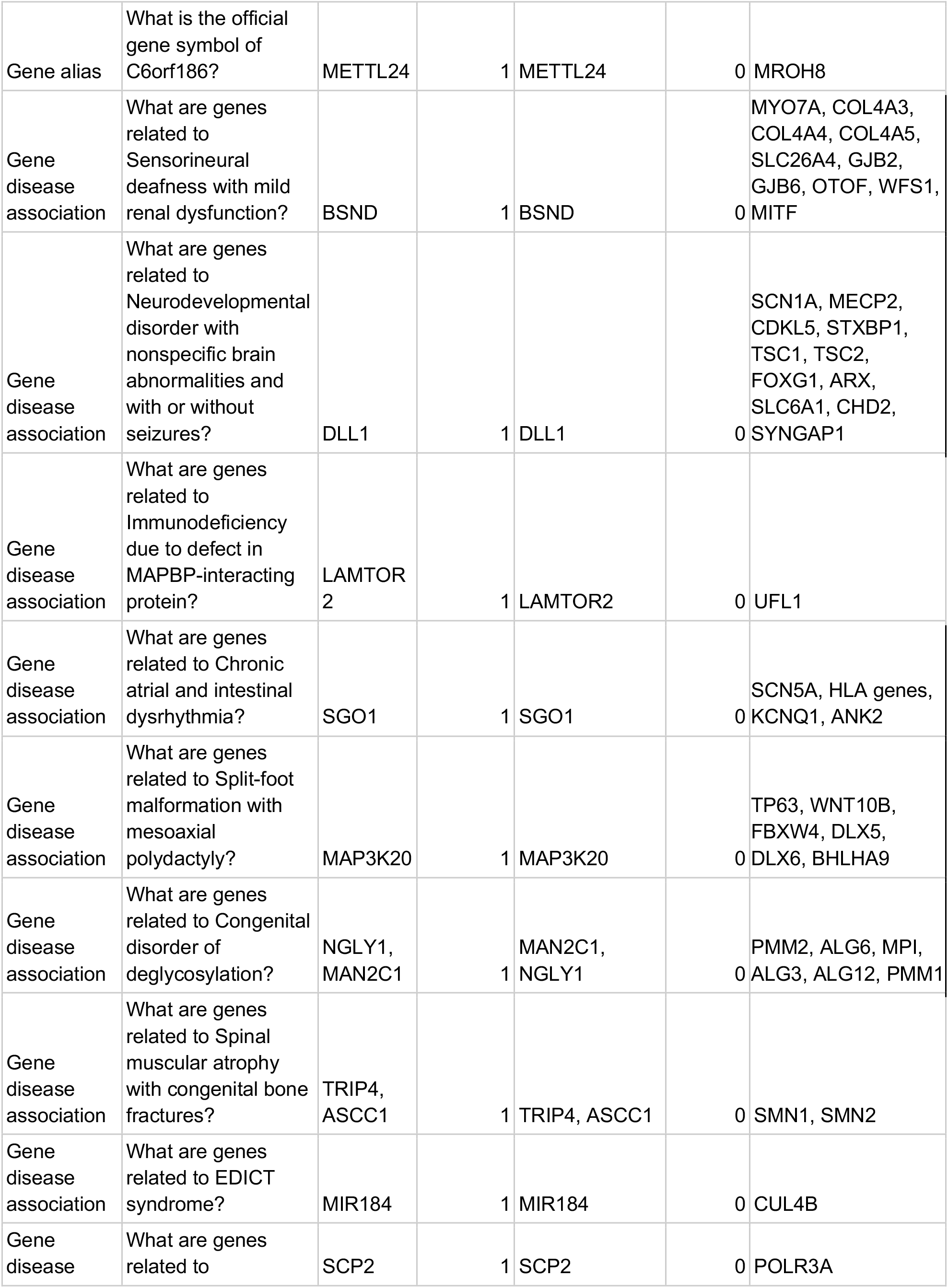
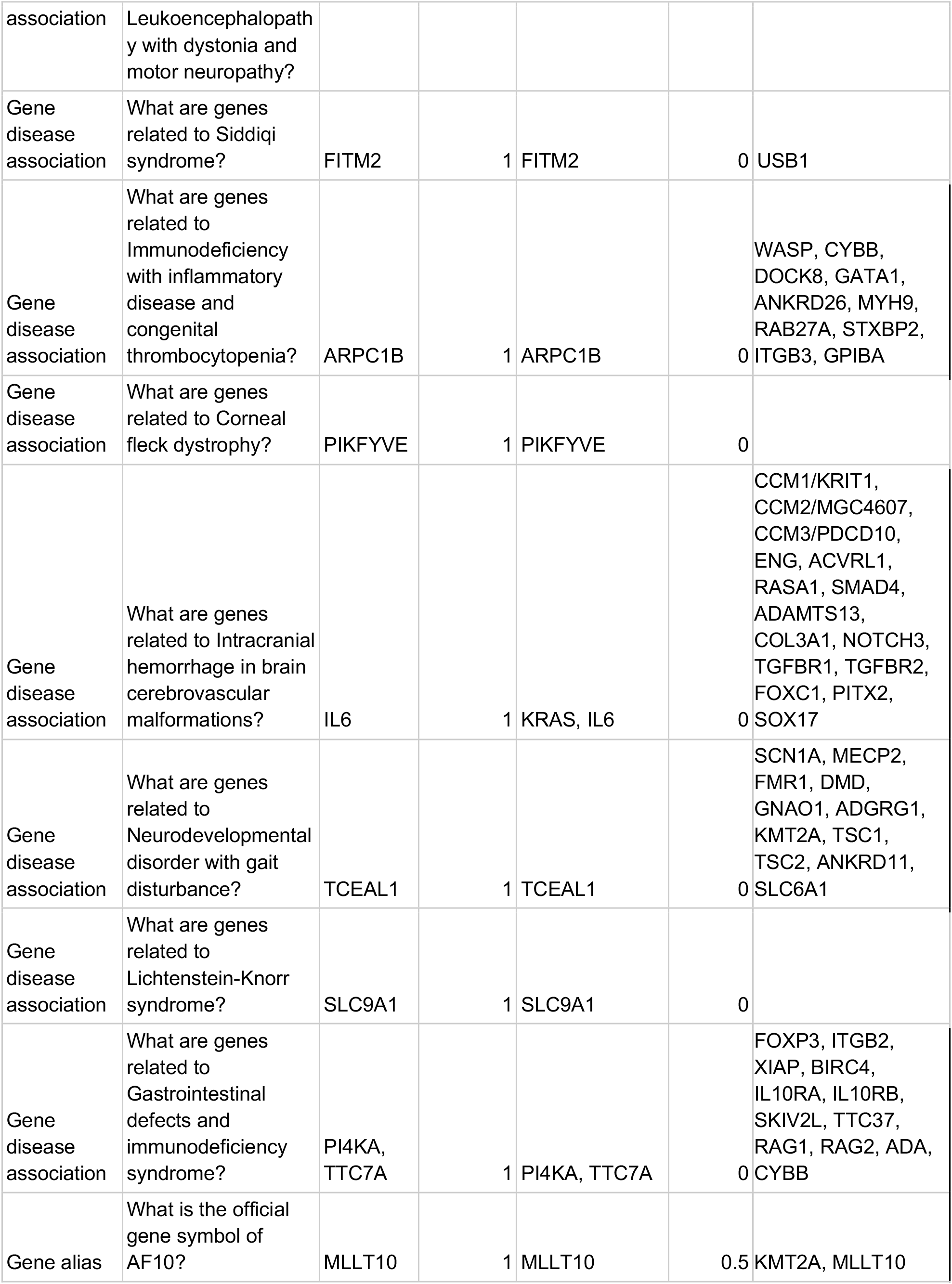
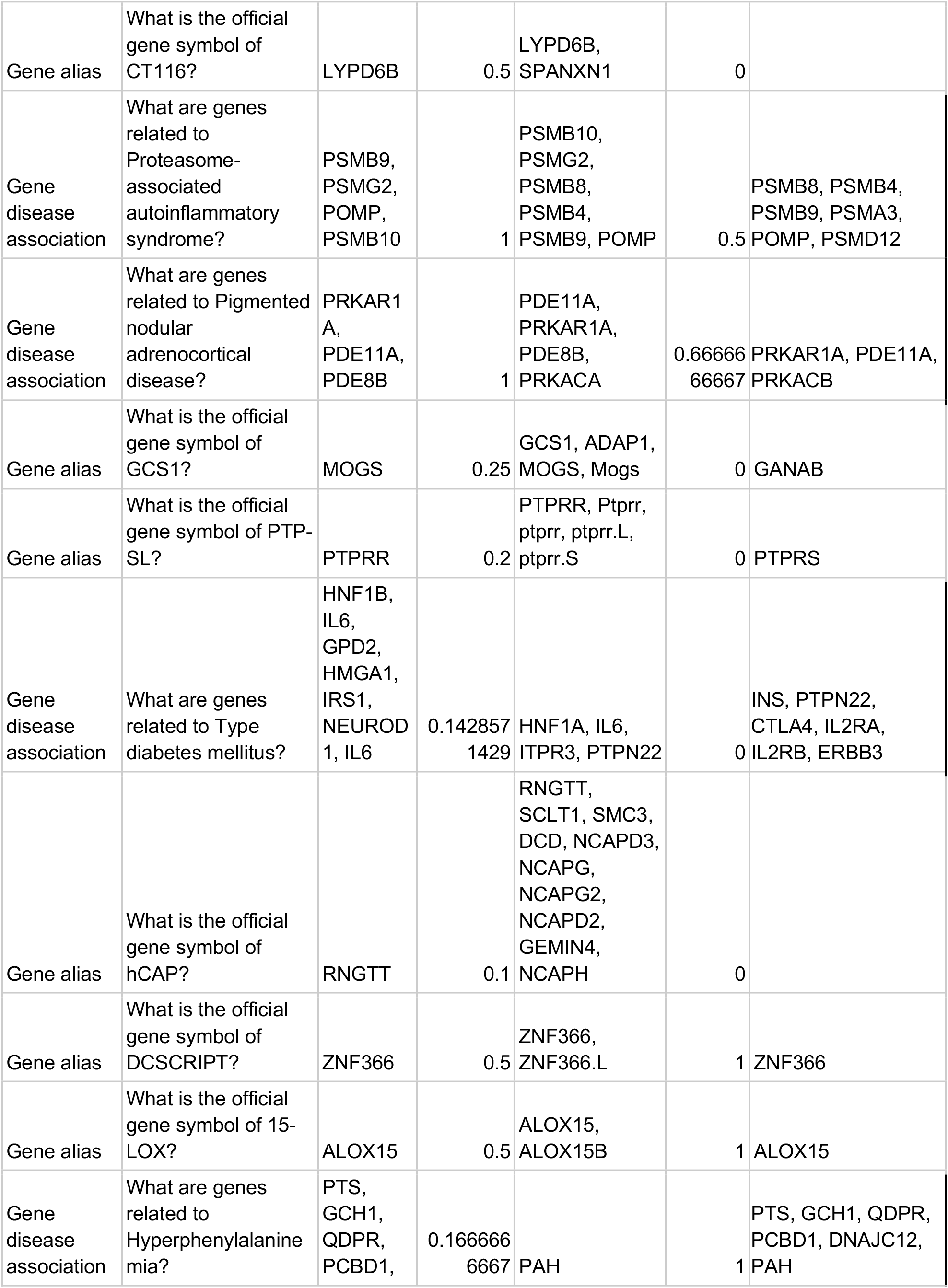
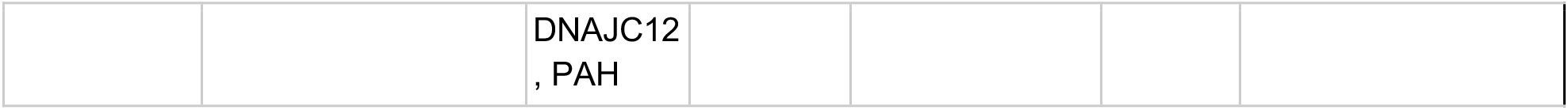
Questions where 4-1106 + Monarch was scored differently than 4-1106 Base, ordered by score difference. Answer Genes columns list gene names extracted from free-text agent answers by evaluator agents. 4-1106 Base without Monarch access outperformed 4-1106 + Monarch in three instances due to scoring criteria and model performance (see Results).

